# Long-lasting coexistence of multiple asexual lineages alongside their sexual counterparts in a fungal plant pathogen

**DOI:** 10.1101/2025.03.28.645883

**Authors:** Ammar Abdalrahem, Axelle Andrieux, Ronan Becheler, Sébastien Duplessis, Pascal Frey, Benoit Marçais, Kadiatou Schiffer-Forsyth, Solenn Stoeckel, Fabien Halkett

**Author notes:** Correspondence: Ammar Abdalrahem, Fabien Halkett.

## Abstract

Deciphering how asexual lineages emerge and persist within otherwise sexual populations is central for understanding the evolution of reproductive modes. In addition to contributing to the long-standing debate about the respective advantages of sexual vs. asexual reproduction, cases of coexistence of both forms also raise ecological questions regarding differences in life-history strategies. Here, we focus on the poplar rust fungus *Melampsora larici-populina,* a plant pathogen species with a complex life cycle and a previously documented case of coexistence of sexual and asexual lineages. We conduct a population genetic study on more than 2,000 individuals sampled over a 30-year period across France to further monitor the persistence of asexual lineages and determine if they form a monophyletic group. Clustering analysis identifies a group of multilocus lineages showing the genetic hallmarks of clonality, including a strongly negative *F*_IS_ value with high variance among loci. This indirect evidence is supported by the repeated resampling of asexual lineages that persist clonally across years. Notably, there is substantial variability within the asexual group, with several genetically distinct multilocus lineages, some of which are entangled with sexual lineages in the phylogenetic tree. The asexual lineages exhibit varying levels of abundance and persistence. One lineage predominates: it has persisted for over 30 years and is more widespread. Asexual lineages co-occur locally with their sexual counterparts, but their prevalence increases southward in France, which is consistent with milder winters being more favourable for asexual overwintering. Together, these results reveal a dynamic and ecologically structured process of repeated switches toward asexual survival with some events of long-lasting persistence. We discuss the consequences of these multiple transitions to asexuality, both in terms of evolutionary ecology and for the management of plant pathogen populations.

## 1 Introduction

The evolution and maintenance of sex remains a fundamental question in evolutionary biology (Culotta & Hines, 1998; Otto, 2009; Hartfield & Keightley, 2012; Neiman et al., 2018). By generating recombinant genotypes through meiosis, sexual reproduction can increase the efficacy of selection, improve the purging of deleterious mutations and enhance adaptation to changing environments (Schurko et al., 2009; Pierre et al., 2022; Parée & Teotónio, 2025). However, it incurs significant costs, such as energetic and time investments required for mate searching (Roughgarden, 1991; Aanen & Hoekstra, 2007; Hörandl, 2013). In contrast, asexual reproduction bypasses these costs by enabling rapid population growth without the need for mates, preserving lineages well-suited to stable local ecological conditions (Shick et al., 1979; Lodé, 2013). Yet, the reduced number of recombination events in asexual lineages reduces the efficacy of selection, rendering them more susceptible to the accumulation of deleterious mutations (Lynch et al., 1993; Peck, 1994; Keightley & Eyre-Walker, 2000), and more likely to lose genotypes through genotypic drift (Rouger et al., 2016; Abdalrahem et al., 2025). The study of species that exhibit both sexual and asexual reproductive modes is therefore valuable for identifying the ecological and evolutionary conditions that favour one reproductive strategy over the other (Butlin, 2002; Simon et al., 2010; Billiard et al., 2012). This is especially relevant in the case of species where asexual lineages repeatedly arise through sexual reproduction events. Here, it is important to note that sexual and asexual reproductions encompass diverse forms, including mixed reproductive modes (Orive & Krueger-Hadfield, 2021), and that the associated costs vary considerably depending on the environmental context, the organisms’ life cycles and life-history strategies (Lehtonen et al., 2012). These factors complement the traditional dichotomy of advantages often attributed to sexual versus asexual reproduction (Butlin, 2002), highlighting the challenge of studying how complex reproductive modes shape evolutionary strategies.

In this study, we focus on a particular type of life cycle in which sexual reproduction occurs once a year and alternates with several episodes of clonal multiplication. Clonal reproduction results in the transmission of the same genotype across generations (with the exception of rare mutations). This life cycle exhibits a two-phase reproductive strategy: during the sexual reproduction phase, dormant forms are produced, enabling persistence through adverse environmental conditions such as winter; conversely, the clonal reproduction phase facilitates rapid population growth during favourable environmental periods. This life cycle is observed in several unrelated species, including aphids (Simon et al., 2002), daphnia (Lehto & Haag, 2010), and fungi (Vialle et al., 2011; Lorrain et al., 2019). In aphids and daphnia, this alternation is commonly referred to as cyclical parthenogenesis (Rouger et al., 2016). Within these species, there exists a polymorphism in reproductive mode. While some lineages follow the full life cycle with an obligate sexual event each year (referred to as *sexual lineages*), other lineages exclusively survive through clonal reproduction for years (Simon et al., 2002; Halkett et al., 2005b; referred to as *asexual lineages*). To avoid any confusion, we use the term ‘clonal’ to refer to the propagation of a genotype during the clonal phase of the cycle (regardless of the type of lineage), and we retain the term ‘asexual’ to describe lineages that survive clonally over the years. Moreover, there is a continuum of investment in sexual reproduction, which is therefore more or less facultative (Halkett et al., 2008). It has also been shown that the level of investment in sexual reproduction correlates with a higher probability that the environment becomes unfavourable, which demonstrates the adaptive nature of these life-history strategies (Halkett et al., 2004, 2008). Examining cases of local co-occurrence of both reproductive modes offer rare opportunities to further decipher the evolutionary ecology of these species, by evaluating the rate of emergence, persistence, and turnover of asexual lineages arising within sexual populations (Lehto & Haag, 2010). However, such ecological and evolutionary histories are far less documented for fungi, which typically lack examples of local co-occurrence of both sexual and asexual lineages.

This study focuses on the phytopathogenic fungus *Melampsora larici-populina*, causing poplar rust, which belongs to the taxonomic order of Pucciniales (Steenackers et al., 1996; Vialle et al., 2011). This species has a complex life cycle analogous to cyclical parthenogenesis, with the additional feature that it switches hosts during sexual reproduction (Figure 1). Thus, two unrelated host plants are required to complete the full life cycle (Hacquard et al., 2011): sexual reproduction takes place in spring on larch trees (*Larix* spp.), whereas clonal (apomictic) multiplication occurs throughout summer on poplar trees (*Populus* spp.), where wind-dispersed spores drive successive epidemic cycles (Pinon & Frey, 2005; Lorrain et al., 2019). In reference to previous publications on aphids and daphnia (Simon et al., 2002), the lineages that complete the full life cycle are therefore hereafter referred to as *sexual lineages*. Sexual reproduction on larch generates recombinant genotypes and obliterates all traces of the previous rounds of clonal multiplication (Xhaard et al., 2011), as theoretically expected under recombination (Aanen & Hoekstra, 2007; Rouger et al., 2016).

**Figure 1.**
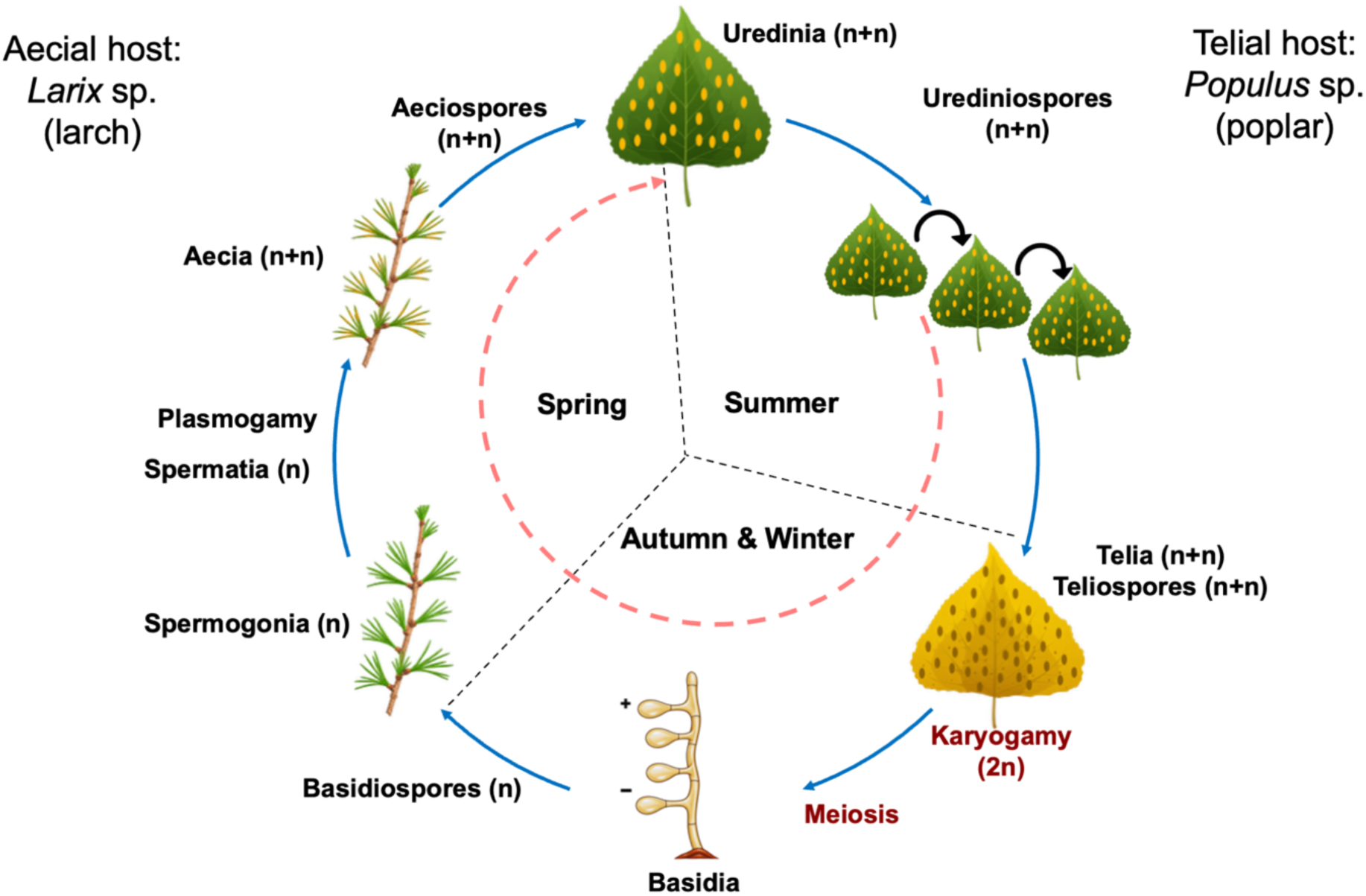
*Melampsora larici-populina* life cycle. In late spring, sexually-derived haploid spores (basidiospores) infect freshly produced larch needles. These infections produce haploid gametes (spermatia) which fuse to generate one (and only one) secondary infection on larch (the aecia). Then *M. larici-populina* mandatorily switch host: the spores produced on larch (aeciospores) are wind-dispersed and can only infect poplar leaves. Note that from the aecial stage onwards, the life cycle is considered diploid, though only cytoplasm fuses, not the two nuclei (dikaryotic cells n+n). Infections on poplar trees result, within 7 to 10 days, in sporulating lesions (uredinia) that initiate repeated cycles of infection and clonal multiplication throughout the season. As a biotrophic fungus, *M. larici-populina* can only multiply on living host tissue. In autumn, the fungus forms frost-resistant overwintering structures (telia). During winter, karyogamy generates transient diploid (2n) nuclei, and meiosis leads to the production of haploid basidiospores that infect larch and complete the life cycle. *Sexual lineages* complete the full life cycle (blue arrows) with mandatory host alternation and an obligate sexual event once a year. *Asexual lineages* can bypass the sexual phase and overwinter through clonal multiplication on poplar alone (inner dashed red arrow). The poplar leaf vector was adapted from “Black Poplar Leaves” (Creazilla), licensed under CC BY 4.0 (https://creazilla.com/media/vector/78070/black-poplar-leaves).

A previous population genetic study (Xhaard et al., 2011) provided evidence for variability in reproductive mode: alongside the sexual lineages, another genetic group displaying all hallmarks of long-lasting clonal reproduction was identified (Halkett, Simon, et al., 2005; Arnaud-Haond et al., 2007; Stoeckel et al., 2021). Signature of long lasting clonal reproduction included a highly negative *F*_IS_ value and significantly over-represented multilocus genotypes (Halkett, Simon, et al., 2005a; Arnaud-Haond et al., 2007; Stoeckel et al., 2021). This suggests that some lineages of the poplar rust fungus could survive through clonal multiplication on poplar alone, without having to necessarily reproduce sexually on a larch tree. In regions where the winter is mild enough, poplars could retain green leaves until very late in the winter, and thus enable the fungus to survive by clonal multiplication until new leaves emerge in spring. This statement coins the ‘*green bridge hypothesis’*, which has never been observed so far and could favour the emergence of *asexual lineages*. Moreover, Xhaard et al. (2011) provided evidence that sexual and asexual lineages can co-occur at the same sampling location. This makes *M. larici-populina* a suitable natural system for studying how asexual lineages emerge and persist within a sexual population. However, this earlier study (Xhaard et al., 2011) was based on only one year of sampling and treated the asexual lineages as a single genetic group. Therefore, it was not possible to confirm that asexual lineages survive clonally over several years, nor to study the number of such asexual lineages implied, their proportion compared to sexual lineages and their genetic diversity characteristics.

Here, we provide an in-depth characterization of the variability in reproductive mode of *M. larici-populina*, with an emphasis on the spatio-temporal persistence of asexual lineages and their genetic characteristics. We further describe the geographic distribution of sexual and asexual lineages and consider the environmental conditions that may favour clonal persistence. Whereas the previous survey (Xhaard et al., 2011) captured the national genetic structure from a single sampling year, the present study adds long-term temporal resampling spanning three decades, including samples collected late in the season to test for the green-bridge hypothesis, and a focused analysis of the emergence and persistence of asexual lineages nested within the sexual gene pool. This study encompasses the analysis of 2,122 individuals sampled across France over 30 years and genotyped using 21 microsatellite markers. Here, we resolve the previously undifferentiated asexual cluster into multiple genetically distinct multilocus lineages (MLLs), and quantify the geographic range, prevalence, and temporal persistence of each. This temporal design allows us to address clonal longevity, repeated emergence, and lineage turnover. In the following, we use a two-fold approach to assess reproductive mode variation among poplar rust individuals. First, we employ an indirect population genetic approach based on clustering among lineages and subsequent evaluation of population genetic indices. Signatures of long-lasting clonal reproduction are: repeated multilocus genotypes (MLGs), linkage disequilibrium, and overall negative *F*_IS_ values with high variance over loci (Balloux et al., 2003; Halkett, Simon, et al., 2005; Stoeckel & Masson, 2014). Second, we take advantage of the temporal sampling to determine whether the same multilocus lineage (MLL) was resampled across years, which provides direct evidence of clonal persistence (i.e. without engagement in the sexual reproduction phase). We then analysed the genetic relatedness among all individuals to resolve the evolutionary relationships among asexual lineages and between them and their sexual counterparts. Finally, we examine local co-occurrence of sexual and asexual lineages, their long-term persistence, and their spatial distribution to gain further insight into the roles of environmental and ecological factors in shaping the survival and maintenance of asexual lineages in this species.

## 2 Material and Methods

### 2.1. Isolation of individuals and sampling strategy

To characterize the temporal and spatial genetic structure of *M. larici-populina* in France, we analysed a total of 2,122 individuals collected from 59 locations (French administrative departments) over three decades (1992-2024) (Figure S1). Each individual originated from a single lesion (uredinium) isolated from a poplar leaf collected *in natura*. The genotypes of all individuals are diploid (sampling during the dikaryotic phase of the life cycle, Figure 1). Note that since individuals were sampled during the phase of clonal multiplication, clonemates (i.e., individuals sharing identical multilocus genotypes except few mutations, having descended exclusively through clonal reproduction since their most recent sexual ancestor) can occur regardless of the lineage overwintering sexually or asexually. To reduce the likelihood of sampling clonemates that would result from local clonal proliferation (Barrès et al., 2012), leaves were sampled on different trees. It is worth pointing out that given the large dispersal ability of this air-borne pathogen (Barrès et al., 2008; Saubin et al., 2024), clone mates belonging to sexual lineages can be found, on rare occasions, in locations separated by around 50 km (Becheler et al. 2017).

The dataset (Table S1) combines previously published genotypes with new additions, structured into three distinct sampling categories:

*(i) National surveys (broad spatial scale):* We integrated data from two major countrywide sampling campaigns. The first corresponds to the 2009 survey (23 locations, n = 428) previously described by Xhaard et al. (2011). The second is a new dataset from a 2011 survey (60 locations, n = 1,374). Individuals from both surveys were collected in late summer (August-September), coinciding with the peak of clonal multiplication prior to the sexual phase (Figure 1). Sampling covered most of the poplar distribution area in France (Figure S1 and Figure 2).
*(ii) Targeted monitoring and historical sampling (temporal scale):* We included 246 individuals from a long-term epidemiological monitoring project in the Durance River valley (2004, 2008, 2020) (Xhaard et al., 2012; Becheler et al., 2016). These were supplemented by 74 individuals from historical samples kept in our laboratory collection, spanning the period from 1992 to 2014.
*(iii) Late-season samplings*: To investigate potential overwintering mechanisms, we sampled from a location in southern France (Cavalaire-sur-Mer) in February of 2014 (n=8) and 2024 (n=40). Unlike poplar stands elsewhere in France, this site retains green foliage throughout the winter, until February. It therefore offers a direct test of the ‘green bridge’ hypothesis, namely whether sporulating lesions can survive on living poplar leaves outside the growing season and thereby allow a lineage to complete its cycle without the larch-dependent sexual phase.

**Figure 2.**
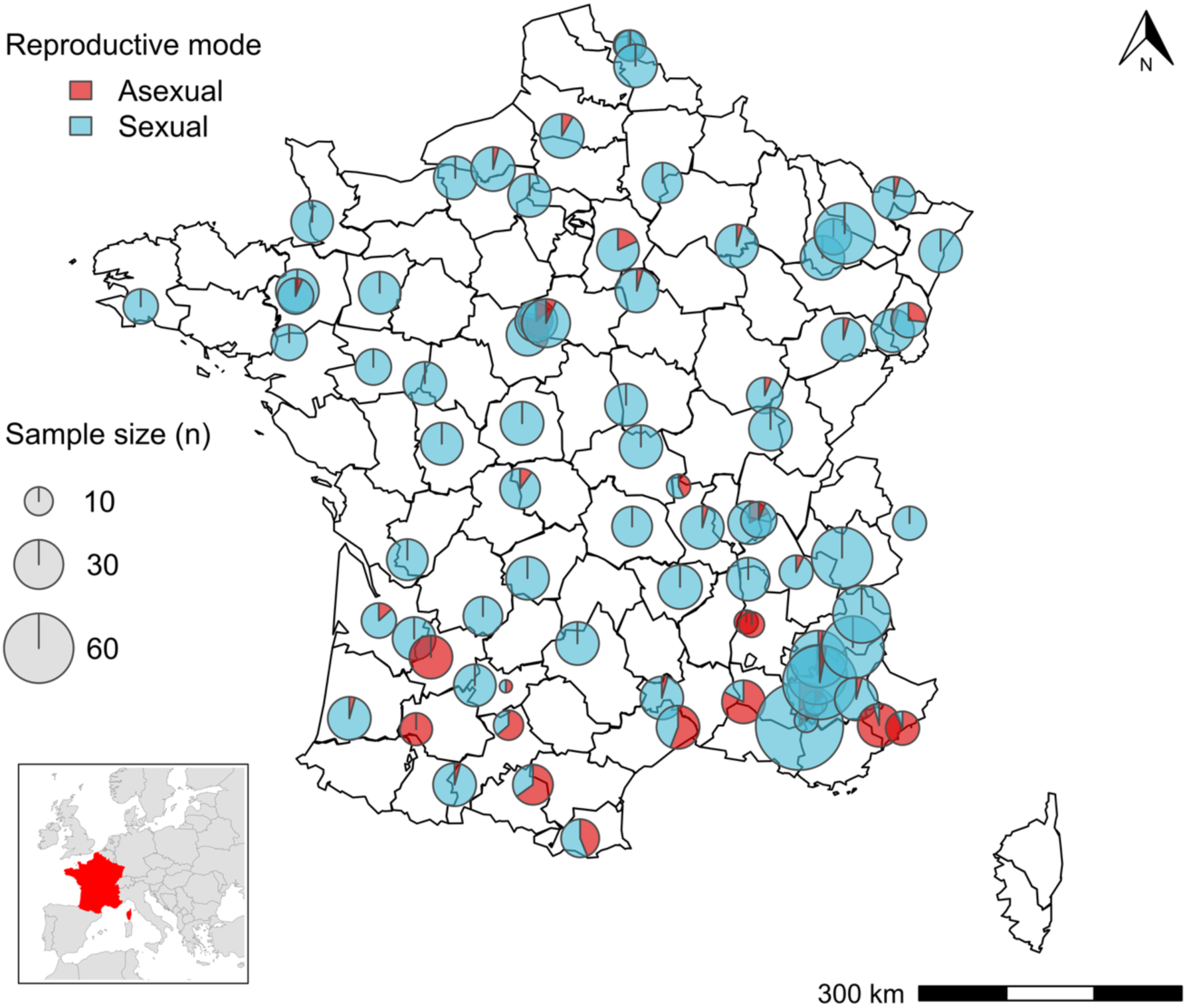
Geographic distribution of sexual and asexual lineages of *M. larici-populina* individuals collected in 2009 and 2011 across France. The pie charts show the proportions of asexual (red) and sexual (blue) lineages at each sampling location. The size of each pie chart corresponds to the total number of individuals collected from that site.

### 2.2 Genotyping

The DNA extraction process was guided by a protocol that employs the BioSprint 96 DNA Plant Kit and the BioSprint 96 automated workstation (Qiagen). Samples were genotyped using 21 microsatellite loci [Mlp_91, Mlp_83, Mlp_50, Mlp_71, Mlp_96, Mlp_95, Mlp_55, Mlp_66, Mlp_12, Mlp_68b Mlp_54, Mlp_57, Mlp_58, Mlp_49, Mlp_93, Mlp_82b Mlp_56, Mlp_87, Mlp_97, Mlp_77, Mlp_94 (Xhaard et al., 2009)]. These loci have been extensively utilized in several population genetic studies of this species (Xhaard et al., 2011; Becheler et al., 2017; Persoons et al., 2017; Louet et al., 2023; Saubin et al., 2024). The amplification through PCR with the Multiplex PCR Kit (Qiagen) was performed as described by Xhaard et al. (2011). It was conducted across three multiplex reactions containing eight, five, and eight loci, ranked by amplification size, respectively. The PCR products were then pooled and analysed using an ABI 3730 Genetic Analyzers (Applied Biosystems) by the company GenoScreen. Fragment sizing utilized a LIZ-1200 size standard, and allele scoring was performed with GENEMAPPER V4.0. Samples that failed to amplify at more than six loci were excluded from further analysis.

### 2.3 Population genetic and statistical analyses

First, to handle the genetic data of microsatellite markers in the R programming language v.4.4.1, we converted the data format from a data frame to a genind object using the df2genind function from the **adegenet** package v.2.1.10. This function optimizes multilocus genotype data and accommodates missing data if it is found in some individuals or loci. Missing data were handled without imputation in the clonal analyses (missing = "asis"). The automated workflow code provides full details on subsequent analyses (Abdalrahem et al., 2026). Moreover, specific analyses were executed out of the main R script: the GenAPoPop 1.0 software was used to assess the discrimination power of the set of microsatellite markers (Waits et al., 2001) and to calculate the *Pareto β* index (Arnaud-Haond et al., 2007; Stoeckel et al. 2021), and the ClonEstiMate software was used to estimate the rate of clonality; Becheler et al., 2017; Stoeckel et al., 2024).

#### Identification of repeated genotypes

The prerequisite to an accurate assessment of reproductive mode is the proper delineation of repeated genotypes derived from clonal multiplication (Halkett, Simon, et al., 2005a; Arnaud-Haond et al., 2007). Individuals with the same combination of alleles (known as multilocus genotypes, MLG) were consistently identified through the R script using mlg.filter from the **poppr** package v.2.9.3 (Kamvar et al., 2014). Genotypes differing by a few alleles were double checked by returning to the original chromatographs, so that all remaining differences were validated. Then we tested the discrimination power of our marker set, to evaluate the probability of finding repeated MLGs by chance (*i.e.* meaning they are not ‘true’ clonemates). To do so, we computed the probability of finding two MLGs that would have been obtained through sexual reproduction, assuming panmixia (*Pidu*) as well as the number of pairs of identical MLGs by chance, given the sample size (*N_Pidu*) using GenAPoPop 1.0 (Evett & Weir, 1998; Waits et al., 2001; Stoeckel et al., 2024).

#### Delineation of multilocus lineages

As the accumulation of mutations over time can lead to slight differences among multilocus genotypes within a given asexual lineage (Arnaud-Haond et al., 2007), we subsequently grouped MLGs into multilocus lineages (MLL). So as not to apply an arbitrary threshold for MLL delineation, we first computed the distribution of pairwise shared allele distance as advised by Arnaud-Haond et al. (2007), and Bailleul et al. (2016). As intra lineage differentiation can vary across MLLs, we chose to use the UPGMA clustering method, which aggregates MLGs one by one by examining the nearest neighbors until the MLL boundaries are found (Kamvar et al., 2014, 2015). The threshold on genetic distance between neighbor genotypes was set using the function cutoff_predictor from **poppr**.

#### Clustering procedure to indirectly identify asexual lineages

Because asexual lineages reproduce without recombination, their genotypes are inherited as fixed multilocus combinations and evolve independently of the recombining sexual population. Over time, this lack of reshuffling causes asexual lineages to diverge genetically from their sexual counterparts. (Rouger et al., 2016; Bast et al., 2018). Indeed, we previously found large genetic differences between sexual and asexual groups of *M. larici-populina* (Xhaard et al., 2011). Clustering individuals would thus be an effective approach to identify asexual lineages without any *a priori*. The main pitfall of this method being that the clustering algorithm should not group individuals according to Hardy-Weinberg expectations, which is the common assumption of any Structure-like approach (Pritchard et al., 2000). We thus carry out our clustering analysis using a PCA and discriminant analysis approach based on allelic distribution (Jombart et al., 2010). We used the find.clusters function from the **adegenet** package to reduce the dimensionality of the dataset. Discriminant analysis of principal components (DAPC) was then used to refine the identification of clusters by maximizing the variance between two clusters, while minimizing within cluster variance (Jombart et al., 2010). Missing alleles for the DAPC were mean-imputed at the allele-frequency level (adegenet::tab, NA.method = "mean"). Cross-validation (xvalDapc; 100 replicates, 90% training set) identified 10 principal components as optimal, with a mean correct reassignment of test-set individuals of 99%, indicating that the two clusters are well separated. These 10 components captured 46% of the total genetic variance. The optimal number of clusters was determined to be K=2, supported by a high Silhouette score in line with our biological hypothesis (Shahapure & Nicholas, 2020), as a statistical support (Figure S2). This clustering aligns with our biological hypothesis about two major reproductive modes (sexual vs. asexual lineages). Accordingly, individuals were assigned using the dapc function with *K*=2. To ensure the reliability of the assignment, only individuals with a high membership coefficient (≥0.8) were retained for further analyses of within-cluster genetic characteristics, and the 7 out of 2,122 individuals (0.3%) that fell below this threshold were excluded from these analyses.

#### Resampling multilocus lineages (MLLs) as direct evidence for asexual reproduction

To track the persistence of MLLs over time, we recorded the years in which each MLL was detected and calculated its frequency across different years. A lineage that persisted for at least two years was considered asexual. We tested the overlap between the direct clustering and indirect resampling approaches for assessing asexual reproductive mode by performing Fisher’s exact test (Clarkson et al., 1993) on the contingency table of MLL sorting according to *both* approaches with the fisher.test function from the **stats** package v.4.4.1.

#### Computation of classical population genetic indices

As the clonal reproductive mode distorts allele distributions within and among individuals, we computed several classical population genetic indices to characterize within cluster and within MLL genetic signatures. Observed heterozygosity (*H*_O_), unbiased expected heterozygosity (*H*_E_) (Nei, 1978), genotype richness (*R*), and multilocus linkage disequilibrium corrected for the number of loci *r̄_d_* were calculated using the **poppr** package (basic function). Allelic richness (*A*_R_) and *F-*statistics, including the inbreeding coefficient (*F*_IS_) (Weir & Cockerham, 1984) were computed using the **hierfstat** package v.0.5-11. The clonal evenness was measured by calculating the *Pareto β* index, which summarizes the distribution in size of MLLs using GenAPoPop 1.0 (Arnaud-Haond et al., 2007; Stoeckel et al., 2024).

#### Spatial analysis of the distribution of sexual and asexual lineages

To illustrate the geographical distribution of sexual and asexual lineages across France, the **maps** package v3.4 was utilized. This enabled us to obtain information not only on spatial distribution, but also on the potential coexistence of the two reproduction modes at a same site. To test the relationship between reproductive mode variation and geographic location, we used a generalized linear mixed model (GLMM) with a binomial distribution and logit link function using the glmer function from the **lme4** package (v. 1.1-35.5) (Bolker et al., 2009). The response variable was the count of sexual and asexual lineages per site. Latitude and longitude were included as fixed effects, and sampling year was included as a random effect. This analysis was performed on the 2009 and 2011 data originating from the two countrywide surveys, so as not to overrepresent certain locations. To further assess the distribution and relative abundance of asexual MLLs across time and space, we performed Fisher’s exact test for fluctuation in MLL abundance across years and locations. Given the large size of the data sets, we employed a Monte Carlo simulation approach with 100,000 replicates to estimate p-values.

#### Genetic relationships between and among sexual and asexual lineages

To further elucidate the evolutionary trajectories of sexual and asexual individuals, and to address the question of the dynamics of transition from sexual to asexual reproduction and vice versa, a discrete matrix based on the relative dissimilarity between individuals was calculated by diss.dist with the **poppr** package to create a neighbour-joining tree (Saitou & Nei, 1987) using nj with 1,000 bootstraps from the **ape** package v5.8.1.

*Inference of the rate of clonality:* Finally, to estimate the rate of clonality in the *M. larici-populina* populations, we employed the ClonEstiMate Bayesian method. This method uses the change of genotype frequencies in a population over several generations to infer reproductive mode (Becheler et al., 2017). We calculated the rate of clonality based on two scenarios: i) merging all individuals within a mixed population (that is any individual can survive asexually by chance), or ii) considering two distinct sexual and asexual populations, based on the lineage sorting detailed above. Since the ClonEstiMate method requires sampled populations to be comparable, we used the data collected during the countrywide surveys carried out in 2009 and 2011.

## 3 Results

### 3.1 Delineation of multilocus lineages

The data set consists of 2,122 individuals genotyped with 21 microsatellite loci. Markers amplified correctly (1930 individuals without missing data, 1.21% missing data overall). We identified 1,470 multilocus genotypes (MLGs) over the 2,122 genotyped individuals. The 21 microsatellite markers we used provided enough statistical power to ensure unambiguous discrimination between genotypes. The overall probability of obtaining the same combination of alleles by chance under panmictic sexual reproduction (*Pidu*) is approximately 10^⁻16^. This implies that 118 million individuals would have to be genotyped before observing by chance two sexually produced descendants with identical genotypes (*N_Pidu*). We thus confidently considered that all repeated genotypes we found were clonemates. We then grouped closely related MLGs into MLLs to account for the few mitotic mutations that could occur during clonal multiplication. The delineation of MLLs was undertaken using a genetic distance threshold of 4.6, as determined by the UPGMA clustering algorithm. This grouping led to the delineation of 1,381 MLLs, so slightly fewer than the number of MLGs, showing that the vast majority of MLGs had no closely related genotype.

### 3.2 Reproductive mode assessment

The clustering approach indicated that almost all individuals exhibited a high probability of membership to one of two genotypic clusters: 450 individuals were unambiguously assigned to cluster one, while 1665 individuals were assigned to cluster two (Figure S3). Only seven isolates had membership coefficients below the threshold and were excluded from further analyses.

A comprehensive population genetic analysis of each cluster (Table 1) was conducted to assess key population genetic indices, revealing a contrast between the two clusters. Cluster 1 exhibits a highly negative *F*_IS_ value, low genotypic diversity, and high linkage disequilibrium (LD), all of which are indicative of a high rate of clonal reproduction (Balloux et al., 2003; Halkett, Simon, et al., 2005a; Arnaud-Haond et al., 2007). The high variance of *F*_IS_ across loci further points to rare events of recombination within a population reproducing essentially asexually over the long term (Stoeckel & Masson, 2014). In contrast, cluster 2 exhibited slightly positive *F*_IS_ values and higher genetic diversity, suggesting a higher degree of allelic recombination, a feature of sexual reproduction. The reduced linkage disequilibrium in this cluster further supported this classification. Consequently, cluster 1 was labelled as asexual, and cluster 2 as sexual. It is noteworthy that similar levels of observed heterozygosity were found in both clusters, but a much reduced allelic richness and gene diversity in the asexual cluster 1. Further, the *Pareto β* index was low (∼ 0.2), indicating a small number of dominant MLGs, which was consistent with high clonality in cluster 1. In the sexual cluster 2, the *Pareto β* was greater than 1, reflecting a more even distribution of genotypes due to genetic recombination.

**Table 1.**
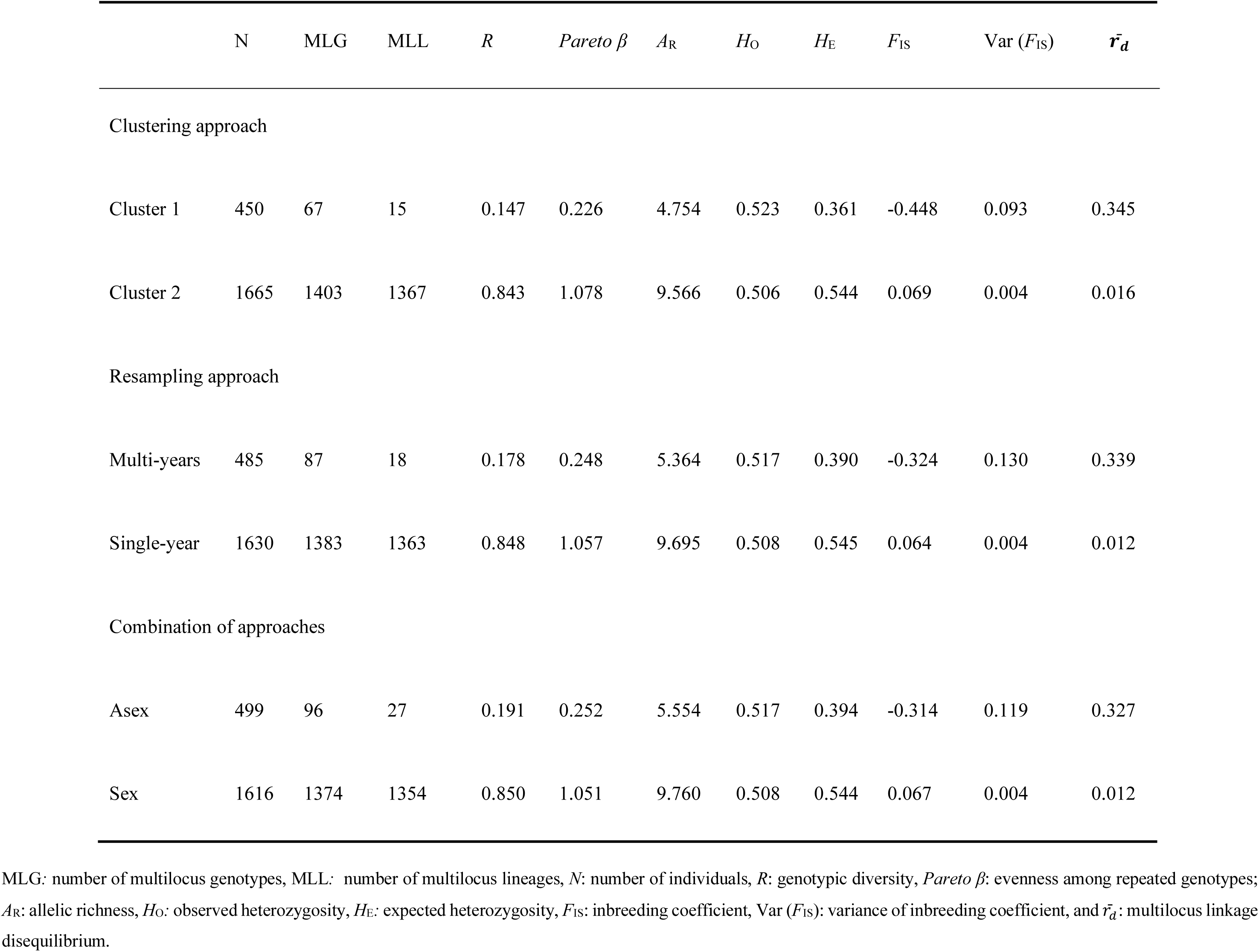
Characteristics of the two genetic clusters identified in M. larici-populina individuals.

|  | N | MLG | MLL | *R* | *Pareto β* | *A*<sub>R</sub> | *H*<sub>O</sub> | *H*<sub>E</sub> | *F*<sub>IS</sub> | Var (*F*<sub>IS</sub>) | ***r̄<sub>d</sub>*** |
|---|---|---|---|---|---|---|---|---|---|---|---|
| Clustering approach |  |  |  |  |  |  |  |  |  |  |  |
| Cluster 1 | 450 | 67 | 15 | 0.147 | 0.226 | 4.754 | 0.523 | 0.361 | -0.448 | 0.093 | 0.345 |
| Cluster 2 | 1665 | 1403 | 1367 | 0.843 | 1.078 | 9.566 | 0.506 | 0.544 | 0.069 | 0.004 | 0.016 |
| Resampling approach |  |  |  |  |  |  |  |  |  |  |  |
| Multi-years | 485 | 87 | 18 | 0.178 | 0.248 | 5.364 | 0.517 | 0.390 | -0.324 | 0.130 | 0.339 |
| Single-year | 1630 | 1383 | 1363 | 0.848 | 1.057 | 9.695 | 0.508 | 0.545 | 0.064 | 0.004 | 0.012 |
| Combination of approaches |  |  |  |  |  |  |  |  |  |  |  |
| Asex | 499 | 96 | 27 | 0.191 | 0.252 | 5.554 | 0.517 | 0.394 | -0.314 | 0.119 | 0.327 |
| Sex | 1616 | 1374 | 1354 | 0.850 | 1.051 | 9.760 | 0.508 | 0.544 | 0.067 | 0.004 | 0.012 |

MLG: number of multilocus genotypes, MLL:  number of multilocus lineages, *N*: number of individuals, *R*: genotypic diversity, *Pareto β*: evenness among repeated genotypes; *A*<sub>R</sub>: allelic richness, *H*<sub>O</sub>: observed heterozygosity, *H*<sub>E</sub>: expected heterozygosity, *F*<sub>IS</sub>: inbreeding coefficient, Var (*F*<sub>IS</sub>): variance of inbreeding coefficient, and *r̄<sub>d</sub>*: multilocus linkage disequilibrium.

The resampling approach revealed two groups: 488 individuals with lineages spanning more than one year due to asexual reproduction and 1,633 individuals ‘not resampled’ through years (likely sexual) (Table 2).

**Table 2.**
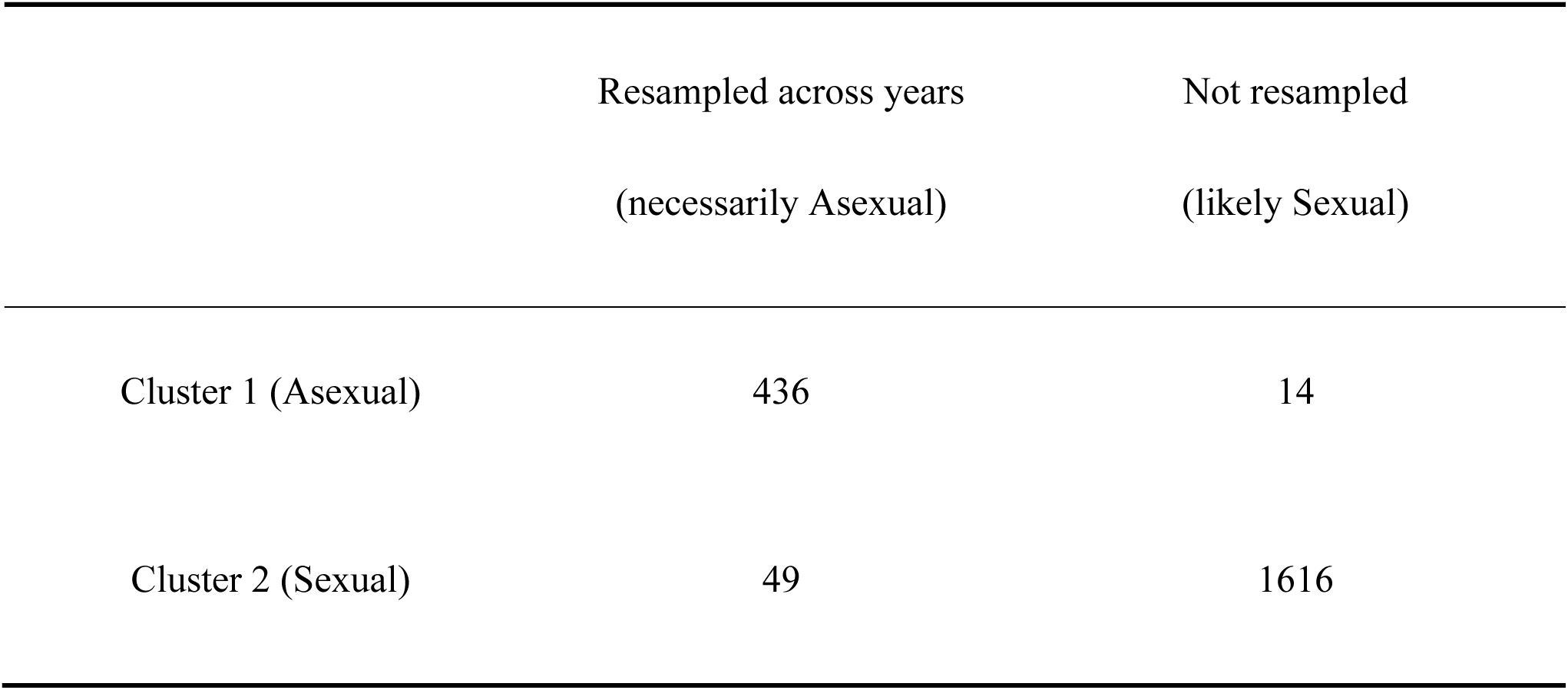
Contingency table showing the relationship between clustering and resampling approaches for reproductive mode definition in the dataset. Values represent the number of individuals assigned to each category.

Comparison of the clustering and resampling approaches showed good agreement between the two methods for defining asexual individuals. A Fisher’s exact test was performed to evaluate the correlation between the assignment of reproductive modes from clustering and resampling analyses. The results were highly significant (p-value < 2.2 × 10⁻¹⁶), thereby rejecting the null hypothesis of no association, with a 95% confidence interval. Only 14 individuals assigned to the asexual cluster were not resampled through years. Conversely around 10% of individuals resampled through years were not assigned to the asexual cluster (Table 2), likely because those asexual lineages emerged too recently to diverge from sexual lineages. In the following, an individual is considered asexual either because its genotype was assigned to the asexual cluster or because the MLL it belongs to was resampled in more than one year. Computing population genetic indices on all asexual individuals showed only slight differences compared to the asexual cluster (Table 1), but some indices revealed a weaker signature of clonal reproduction, suggesting that individuals resampled through years but not assigned to the asexual cluster emerged more recently.

### 3.3 Spatial distribution of reproductive modes

The spatial distribution of both sexual and asexual lineages of *M. larici-populina* individuals in France revealed distinct patterns (Figure 2). Sexual lineages were widespread across most regions, predominating in the northern and central parts of the country. In contrast, asexual lineages were more concentrated in the southern regions.

To further investigate the factors that influence the prevalence of asexual lineages, a GLMM binomial regression analysis was conducted. The analysis (Table 3) revealed a significant effect of both latitude and longitude (Figure S4) on the proportion of sexual and asexual lineages, with positive coefficients indicating that the relative abundance of asexual individuals increases with lower latitude (southwards). Additionally, the random effect of years (variance = 0.12, SD = 0.35) indicated that the temporal distribution of individuals was not associated with a specific reproductive mode.

**Table 3.** Results of the binomial regression model predicting the effect of latitude and longitude on the reproductive mode of *M. larici-populina*.

| Predictor | Estimate | Std. Error | z-value | p-value |
|---|---|---|---|---|
| (Intercept) | - 19.15 | 2.15 | -8.87 | < 2e-16 |
| Latitude | 0.45 | 0.04 | 9.61 | < 2e-16 |
| Longitude | 0.08 | 0.02 | 3.09 | 0.001 |

Estimate (regression coefficient), Std. Error (standard error of the estimation), z-value (z-statistic), p-value (statistical significance)

### 3.4 Inference of clonal rate

The ClonEstiMate Bayesian analysis yielded insights into the rate of asexual reproduction in *M. larici-populina*. The 99% credible interval from the posterior probability was obtained for each group’s clonal rates. In the context of a mixed group, where sexual and asexual lineages are merged into one population, the estimated clonal rate was between 0.31 and 0.38. When the analysis was performed under the assumption of two distinct populations, the clonal rate was between 0.8 and 0.9 for the asexual group and between 0.12 and 0.2 for the sexual group.

### 3.5 Genetic relationships between sexual and asexual lineages

The dendrogram tree shows different clades for sexual and asexual lineages (Figure 3), highlighting seven main asexual lineages. However, some asexual lineages also appeared as closely related sister branches to the sexual clade. Barring MLL9, one of the most abundant asexual lineages, all of the asexual MLLs closely related to the sexual clade are of small size. These small asexual lineages were thus sporadic, unlike the seven most abundant asexual MLLs that show persistence through time (Figure 3, Table 4). These main asexual MLLs also displayed some reticulation, indicating they consist of different MLGs. In contrast, MLL5 (a large MLL classified as sexual) consists of individuals sampled only in 2024 that have exactly the same genotype. These observations underscored the evolutionary relationships and temporal dynamics between reproductive modes within and among *M. larici-populina* lineages.

**Table 4.**
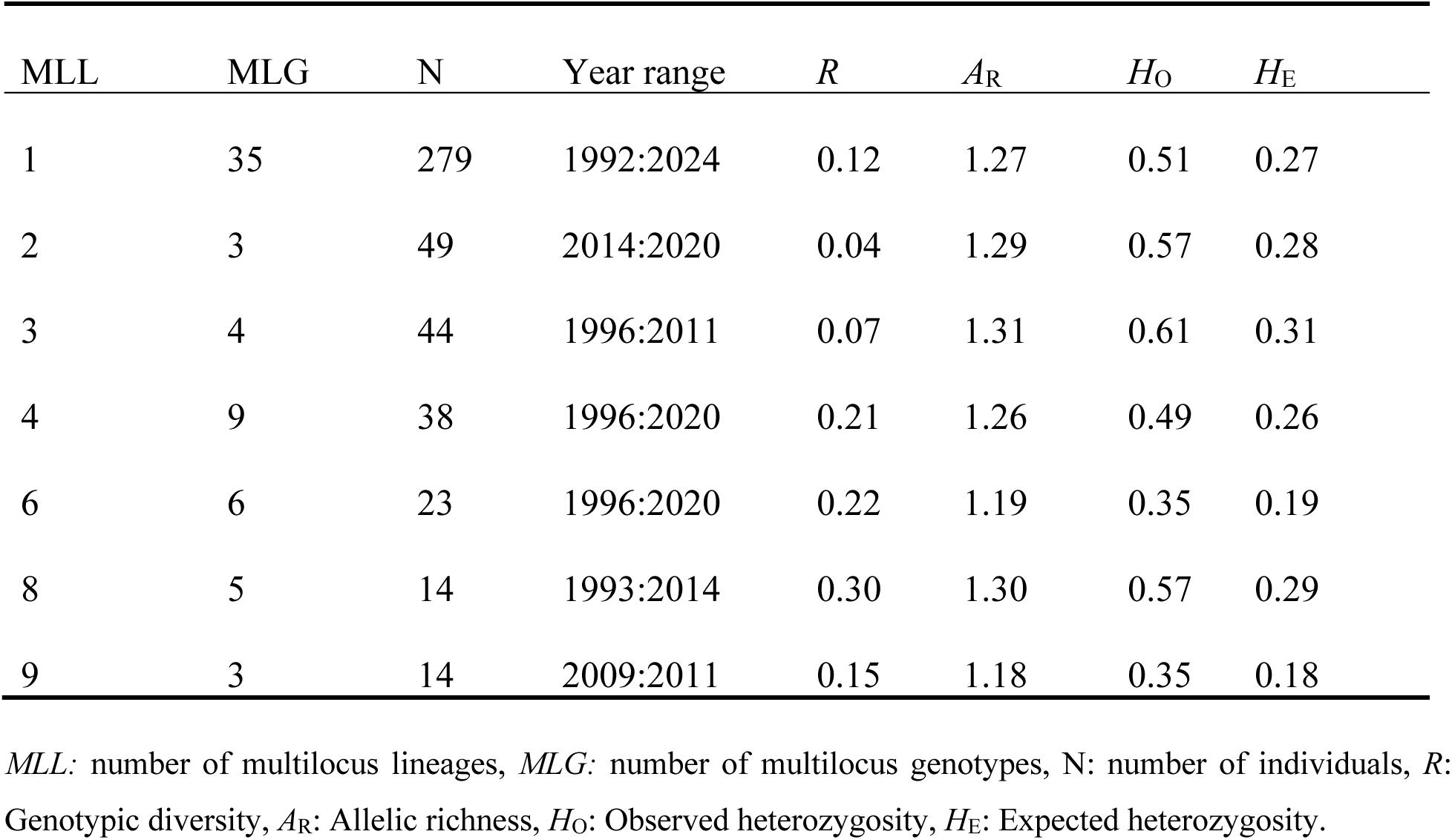
Characteristics of the seven most abundant asexual multilocus lineages (MLLs) of *M. larici-populina*.

**Figure 3.**
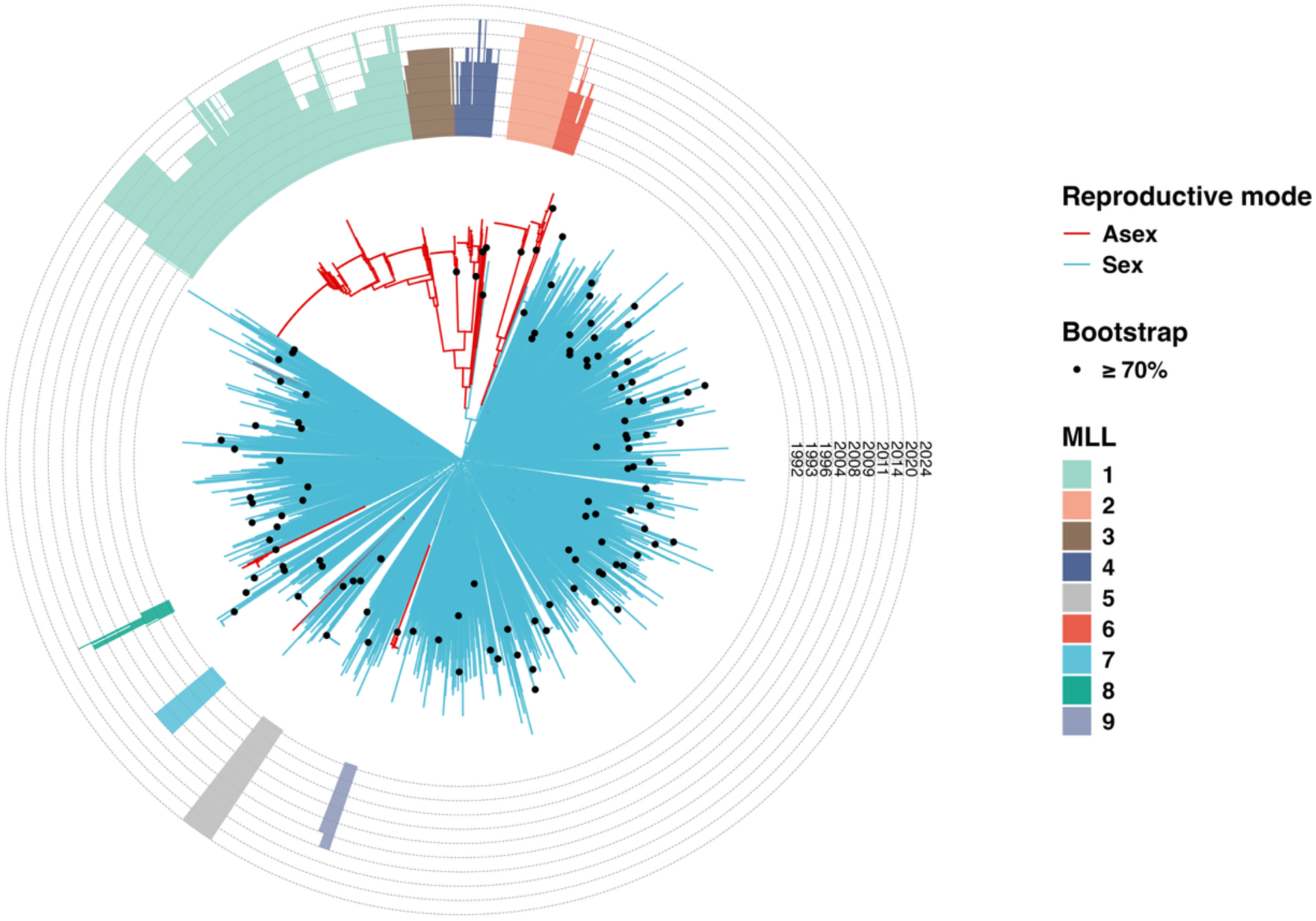
Circular Neighbour-joining (NJ) tree based on microsatellite data (21 markers) illustrating the genetic relationships among *M. larici-populina* individuals. The colour of the branches indicates the reproductive mode, with blue representing sexual and red representing asexual modes. The outer rings represent the sampling year, with each height corresponding to a year. The nine most abundant multilocus lineages (MLLs) are color-coded.

### 3.6 Genetic diversity and persistence of asexual lineages

A more detailed analysis of the asexual group identified several distinct genetic lineages. Among these, seven lineages were found to be the most abundant. These lineages exhibited varying levels of persistence over the years, as indicated by their range of collection years (Table 4). In particular, MLL1 was consistently detected in almost all collection years and across all regions, highlighting its long-term persistence and widespread occurrence.

Within the same lineage (MLL), most variants differ from the main genotype at a single locus (60%) or two loci (22%). It should be noted that, most of the time, the allele difference consists in a change of one repeat motif. Variants differing at three or more loci, as expected from recombination, are rare (17% of variants). This agrees with the low rate of sexual reproduction estimated for these lineages (ClonEstiMate, c ≈ 0.93–0.99), confirming that rare sex may still occur, but mutation is the main source of within-lineage diversification.

The distribution of these lineages in France differed with some lineages being widespread and others more restricted to some areas (Figure 4).

**Figure 4.**
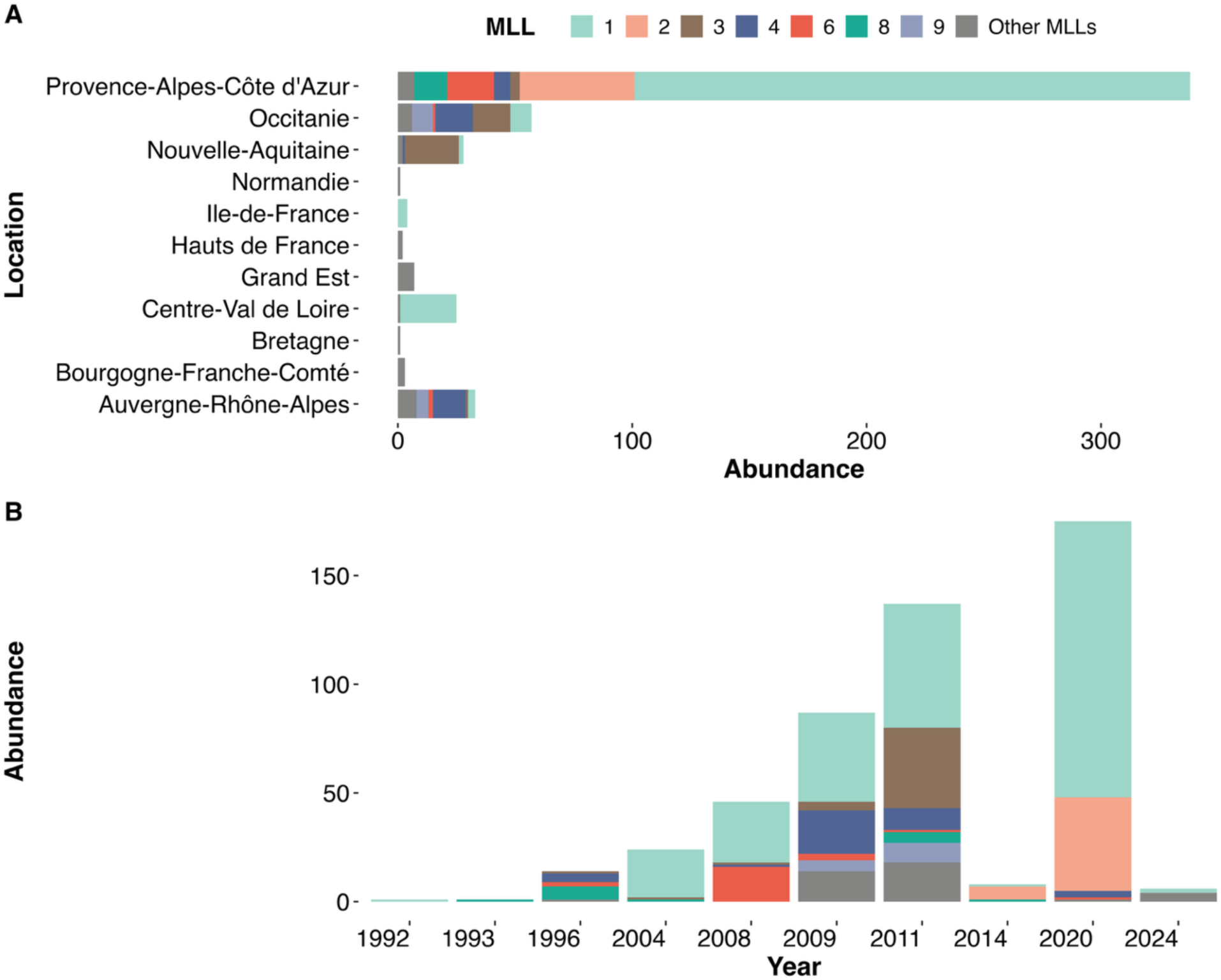
Distribution of asexual lineages over time and across locations. (A) The abundance of asexual MLLs in different French administrative regions. (B) The abundance of asexual MLLs in different years.

Fisher’s exact test revealed a highly significant association between MLL distribution and both temporal and spatial factors (P = 1e-05). The analysis revealed that certain MLLs were repeatedly observed in the same geographic regions and over multiple sampling years, suggesting a pattern of persistence. Conversely, other MLLs appeared sporadic.

### 3.7 Specific analysis of samplings in the late season

The detailed analysis of the samples collected at Cavalaire-sur-Mer in February gave discrepant results depending on the year of sampling (Table S2). In 2014, only eight individuals were analysed. All were assigned to the asexual group and belonged to three lineages. One individual belonged to the predominant asexual lineage (MLL1); another belonged to MLL8 sampled from 1993 to 2011 but not thereafter, and the remaining six belonged to MLL2 that was extensively resampled in 2020 at several locations in the Durance river valley, but never before that. In 2024, out of the 40 genotyped individuals, only six belonged to the asexual group (two belonged to MLL1 and four belonged to MLL35 assigned as asexual but never resampled). Two-thirds of the individuals were clonemates and belonged to MLL5 (see Fig. 3), a lineage assigned to the sexual group and never resampled. The remaining 12 individuals are members of two MLLs sampled three and two times respectively, and three unique genotypes.

## 4 Discussion

The poplar rust fungus *M. larici-populina* is commonly regarded as an obligate sexual species that needs the presence of both poplars and larches to perform its annual life cycle (Pinon & Frey, 2005; Hacquard et al., 2011). However, a previous population genetic study suggested the existence of a distinct group of individuals that would survive asexually only on poplar hosts (Xhaard et al., 2011). Here i) we characterized the existence of an asexual group persisting on the poplar hosts over the years, ii) we found that this asexual group is made up of several genetic lineages. We documented that some of these asexual lineages have persisted for nearly three decades despite their coexistence with sexual lineages in some places. These findings challenge conventional assumptions about the evolutionary dominance of sexual recombination in this species and offer a unique opportunity to investigate the transitions from sexual to asexual reproduction in a plant pathogenic fungus.

In the present study, two complementary approaches were employed to reveal and characterize asexual individuals. A clustering approach based on differences in allelic frequency and probability of membership was combined with a resampling approach aiming to identify the persistence of lineages over the years. We observed a strong overlap between these indirect and direct approaches, which enabled a sound delineation of asexual lineages among *M. larici-populina* individuals. This result further supports the reliability of the clustering approach performed in the initial study of Xhaard et al. (2011) to identify asexual lineages indirectly. Applying such an individual based clustering approach is all the more relevant because sexual and asexual lineages can be admixed in the same location (Halkett et al., 2006). In addition to the genetic differentiation between sexual and asexual clusters caused by the lack of gene flow, several genetic signatures clearly distinguished asexual lineages, thereby reflecting the fundamental differences in evolutionary trajectories between these reproductive modes (Balloux et al., 2003; Halkett et al., 2005a; Rouger et al., 2016). Sexual reproduction ensures the convergence toward Hardy-Weinberg equilibrium, with recombination among loci that reduces linkage disequilibrium. Asexual reproduction, on the other hand, distorts both allelic and genotypic frequencies mostly because genotypes are transmitted clonally as intact multilocus combinations rather than being reshuffled by recombination and segregation (Stoeckel & Masson, 2014; Rouger et al., 2016). As a result, the best way to identify asexual lineages is to apply a multi-criteria method (Kronauer et al., 2012) in which the (diploid) population displays the following hallmarks (Halkett, Simon, et al., 2005; Arnaud-Haond et al., 2007): a reduced genotypic diversity, combined with a significant excess of heterozygotes (negative *F*_IS_ values). This excess of heterozygotes has long been attributed to the independent accumulation of mutations on the two chromosomes (Meselson effect), but recent theoretical analyses suggest that the main evolutionary force causing the fixation of heterozygotes is genetic drift, as it more heavily impacts clonal populations (Stoeckel & Masson, 2014; Reichel et al., 2016; Abdalrahem et al., 2025). It should be noted that we also observed a high variance of *F*_IS_ values across loci, as expected when recombination events seldom occur (Balloux et al., 2003; Stoeckel & Masson, 2014; Stoeckel et al., 2021). The asexual group also displayed significant linkage disequilibrium among loci, which is often reported in asexual and partially asexual species, despite not being consistently expected in neutrally evolving populations (Stoeckel et al., 2021). Linkage disequilibrium was also observed in the sexual group along with a positive *F*_IS_ value as the result of a Wahlund effect. Indeed, the sexual group encompasses different genetic subgroups that have been identified by previous population genetic studies (Xhaard et al., 2011; Persoons et al., 2017) and which are not presented here. We observed clonemates in the sexual group too, as these can form lineages as abundant as those in the asexual group, with the notable difference that they always occur at a single sampling site, and critically that they are never resampled over the years. The local abundance of clonemate lineages is explained by the sampling being done at the end of the clonal epidemic phase of the *M. larici-populina* life cycle. It should be recalled that the sexual reproduction obligately takes place on larch and resets the population genetic indices in the sexual group each year (Xhaard et al., 2011; Persoons et al., 2017). The contrast in population genetic indices provides a sound framework for distinguishing reproductive modes, supporting the reliable assignment of individuals into either sexual or asexual groups and providing the first insights into their respective evolutionary history (Xhaard et al., 2011).

Asexual lineages have often been described as distributed at the margins of a species’ natural range (Simon et al., 1999; Halkett et al., 2005a; Drenth et al., 2019; Pereyra et al., 2023), a phenomenon called ‘geographic parthenogenesis’ (Peck et al., 1998). Similarly, invasive species often shift towards asexual reproduction in their novel range, while reproducing sexually in their native environment (Song et al., 2011; Platt & Jeschke, 2014; Gladieux et al., 2015; Tilquin & Kokko, 2016). For example, asexual reproduction is dominant in invasive populations of some ant species, such as *Wasmannia auropunctata* and *Cerapachys biroi* (Foucaud et al., 2006; Kronauer et al., 2012). Plant pathogenic fungi also adopt diverse reproductive strategies, with asexual reproduction being responsible for the most damaging epidemics (Ashu & Xu, 2015; Gladieux et al., 2015; Drenth et al., 2019). Potato late blight is a devastating disease caused by the oomycete pathogen *Phytophthora infestans.* Both sexual and asexual reproduction take place in the native range, but it is asexual reproduction only that led to the devastating Irish Potato Famine in the 1840s (Gavino et al., 2000; Ristaino, 2002; Fry, 2020). The wheat yellow rust fungus *Puccinia striiformis* f. sp. *tritici* displays a similar pattern, with different reproductive modes depending on the region. The fungus reproduces sexually in China and Pakistan, which is highlighted by its high genotypic diversity, whereas it reproduces asexually in the rest of the world (Bahri et al., 2009; Mboup et al., 2009; Ali et al., 2014). The rice blast fungus *Magnaporthe oryzae* (Syn: *Pyricularia oryzae*) reproduces sexually in a limited area of Southeast Asia, its native range, whereas epidemics globally are driven by asexual lineages (Saleh et al., 2012). Here, in accordance with the previous study of Xhaard et al. (2011), we found that the asexual group was unevenly distributed across the country and mostly occurred in southern France. Poplar rust is intimately dependent upon the presence of its two hosts. Although the native range of European Larch (*Larix decidua*) is restricted to the Alps and some other Central European mountain ranges, it has been widely planted in many parts of France in mountains and plains since the 18th century. Nevertheless, it is less frequent in the Mediterranean area, which can therefore be considered as the margin of the poplar rust fungus’ range. If we compare our results with those of other rust fungi, particularly those infecting wheat, we can see a larger spatial segregation between the sexual lineages found only in native distribution area of the pathogen species and the asexual lineages that have invaded the crops worldwide. Interestingly, asexual lineages in rusts affecting food crops are specialized on different cultivars, and therefore on different host genotypes (Goyeau et al., 2007). The relative abundance of asexual lineages therefore fluctuates over time with the use or disuse of these cultivars (Fontyn et al., 2022). This shows to which extent the asexual evolution of these plant pathogen species is driven by human practices. By contrast, in *M. larici-populina*, asexual lineages are sampled mostly on wild poplar trees and are thus found on different host genotypes. We thus can argue that this rust species enables the incipient stage of sexual to asexual transition to be documented.

A puzzling observation is that asexual lineages of *M. larici-populina* co-occur with their sexual counterparts in most sampling sites. The absence of larch trees in some areas can be compensated for by the very high dispersal abilities of this species (Barrès et al., 2008; Saubin et al., 2024), and explains why sexual and asexual lineages are admixed even if no larch tree exists in the close vicinity of the sampling site. Another non-exclusive hypothesis to account for the long-lasting coexistence of sexual and asexual lineages is that of a temporal fluctuation in the environmental conditions that favour one or the other. Indeed, almost half the asexual lineages were sampled in 2020 alone. These fluctuations in relative sexual vs. asexual abundances could be linked to particular meteorological events that desynchronize the alternation between poplar and larch during the sexual life cycle. This could cause a disadvantage to sexual lineages that have to complete this complex life cycle. An outstanding example for the effect of a hazardous event on the fate of sexual and asexual lineages was documented in the red alga *Agarophyton chilense* along the Chilean coasts (Becheler et al., 2020). This study monitored wild (mostly sexual) and cultivated (asexually propagated) populations of red alga before and after a devastating earthquake. Although sexual populations paid a large tribute immediately after the catastrophe, the artificial maintenance of an asexual population caused the algae culture to collapse in the long term, suggesting that each mode of reproduction plays a role in maintaining a dynamic equilibrium.

The observed distribution of asexual lineages of *M. larici-populina,* with a higher prevalence in southern France, points to a mechanism that could explain their maintenance and diversification. In milder climates the asexual survival could be facilitated by a longer growing period of poplars, creating a ‘green bridge’ that would enable the rust lineages to jump over the winter from poplar leaves to poplar leaves. According to this hypothesis, the asexual lineages would passively bypass the sexual phase on larch, without the need for any genetic determinism. In support of this hypothesis, it should be noted that *M. larici-populina* urediniospores can survive for 10 months and still be able to infect poplar leaves (Taris, 1968). But in these experiments, rust-infected poplar leaves were stored either in a greenhouse, or in a wire cage 2 meters above ground level, which does not mimic real natural overwintering conditions. According to this neutral hypothesis, both the advent and the survival of asexual lineages would be a matter of chance only. In support of this hypothesis, we observed a large lineage (MLL5) assigned to the sexual group and sampled in winter (February) in Cavalaire-sur-Mer. This is the direct observation that a sexual lineage can bridge over the winter on poplar leaves. In addition, two asexual lineages (MLL8 and MLL9) branched among the sexual cluster and were considered as asexual solely because they were resampled through years. These observations document the incipient stage of the emergence of asexual lineages.

The persistence of several asexual lineages over decades, attesting to their evolutionary maintenance and relative success, could also be indicative of some genetic determinism favouring their survival over winter. In support of this hypothesis, the trade-off between overwintering ability and multiplication during the epidemic season has been documented in other fungal species (Hamelin et al., 2016; Suffert et al., 2018), and proven to account for the coexistence and evolutionary maintenance of sexual and asexual lineages (Castel et al., 2014). It is also likely that long-lasting asexual lineages could have lost, at least to some degree, the ability to invest in sexual reproduction, maybe to the benefit of greater asexual survival. Our population genetic analysis, and especially the high variance of *F*_IS_, would nonetheless indicate that recombination does not occur among asexual lineages and thus that they do not evolve under strict asexuality but rather as a succession of clones maintaining on poplar leaves over winter, renewed over the years.

Furthermore, the rare engagement of asexual lineages into the sexual reproduction can bring additional explanations for the diversity of asexual lineages we observed. In line with the concept of contagious asexuality observed in other organisms with mixed reproductive modes, such as aphids and daphnia (Simon et al., 2002, 2003; Halkett et al., 2006, 2008), asexual lineages that invest in sexual reproduction could be able to produce offspring that inherit their propensity to survive asexually (Halkett et al., 2008). Further deciphering the history and evolutionary success of these asexual lineages calls for a dedicated experiment to assess their performance and investment throughout the life cycle (Pernaci et al., 2014; Abdalrahem, 2025).

The main original result of our study was to document the diversity among asexual lineages in *M. larici-populina*, with some lineages persisting over decades, and others appearing more sporadically. These patterns are best understood as an emergence-persistence continuum rather than a single, ancient origin of asexuality. Asexual lineages seem to arise repeatedly and independently from the sexual population, with the most recently emerged lineages being polyphyletic and nested within the sexual gene pool. Only a few lineages, MLL1 being the most obvious, become established over the long term and across wide areas, while most remain local or short-lived. The factors that distinguish successful from short-lived lineages probably combine the opportunity for asexual overwintering, favoured by the milder winters of southern France, the founder genotype and its fitness, and the local host and environmental conditions. The fact that two sampling campaigns covered the entire French territory, and that other sites were sampled regularly over more than 30 years supports that this pattern is not due to sampling bias. We documented that asexual MLLs can consist of different genotypes (MLGs), with some long-lasting asexual lineages such as MLL1 showing relatively high genotypic diversity. Conversely, lineages within the sexual group primarily consist of a singular repeated MLG. Here, it is important to reiterate that MLGs within asexual MLLs differ only slightly; in the vast majority of cases, the differences are limited to no more than three loci. Moreover, the changes consist in either heterozygous to homozygous conversion or an increase or decrease in allele length by a one-step mutation. This regular pattern of allele changes, found across all MLGs within MLLs, supports the hypothesis that the asexual lineages evolved through single mutation steps. Although mutation rates for the microsatellite markers used in *M. larici-populina* have not yet been estimated, this regular pattern could present an opportunity to apply a tip calibration approach of molecular clock analysis to estimate the timing of the emergence and persistence of asexual lineages (Rieux & Balloux, 2016). Yet, the differential abundance of MLGs and genotypic diversity among asexual lineages reflects heterogeneity within asexual population dynamics and emphasizes the necessity to consider each lineage independently, rather than assuming that the whole asexual population follows the same evolutionary trajectory. The most parsimonious hypothesis is that this diversity has been generated by the repeated emergence of asexual lineages through the evolutionary mechanisms previously discussed (chance of survival or contagious asexuality). Among all asexual lineages, MLL1 stood out as the most abundant lineage, being repeatedly sampled from 1992 onwards and showing an emergence in 2020. It was detected in all regions where asexual lineages were sampled, and forms the only contribution to the unexpected occurrence of asexual lineages northward in France, nearby Paris. The genome of a reference asexual individual, representative of this MLL, has recently been sequenced, and analysis of its genome structure, polymorphisms, and gene content will further document the tale of this dominant asexual lineage.

## Data

To ensure reproducibility, all analyses were executed using R version 4.4.1. The relevant scripts and data can be accessed at https://doi.org/10.5281/zenodo.15100450 (Abdalrahem et al., 2026).

## Author Contributions

Sébastien Duplessis, Pascal Frey, and Fabien Halkett conceived and designed the study. Axelle Andrieux designed and performed most of the molecular experiments. Ronan Becheler, Ammar Abdalrahem, Axelle Andrieux, and Fabien Halkett performed the manual curation of genotyping results. Axelle Andrieux, Ronan Becheler, and Fabien Halkett did the preliminary analyses of the data. Ammar Abdalrahem performed the final data analyses with help and advice from Benoit Marçais, Ronan Becheler, Solenn Stoeckel, and Fabien Halkett. Ammar Abdalrahem, Kadiatou Schiffer-Forsyth, and Fabien Halkett prepared the manuscript. All authors revised and approved the manuscript.

## Acknowledgments

We thank Jean Pinon for having created the historical poplar rust collection at INRAE Nancy in the 1980s and for having sampled the Cavalaire-sur-Mer population in 2014 and 2024. We warmly thank Asmaa A. Eltahlawy and Scott Heslop for proofreading an earlier version of the manuscript. This long-term research program was supported by several grants from the French National Research Agency (ANR-11-BSV7-0007, CLONIX project; ANR-12-ADAP-0009, GANDALF project; ANR-13-BSV7-0011, FUNFIT project; ANR-18-CE32-0001, CLONIX2D project; ANR-23-CE20-0032, ENDURANCE project). This research program was part of the Arbre LabEX (ANR-11-LABX-0002-01) and the interdisciplinary program ARTEMIS of Lorraine Université d’Excellence (ANR-15-IDEX-04-LUE). Ronan Becheler was supported by a postdoctoral fellowship from both the Region Lorraine and the French National Research Agency (CLONIX project, ANR-11-BSV7-007). Kadiatou Schiffer-Forsyth is supported by a PhD fellowship from both the Region Grand Est and the French National Research Agency (ANR-23-CE20-0032, ENDURANCE project). Ammar Abdalrahem was supported by a PhD fellowship from the French Ministry of Education and Research (MESR).

We have used AI tools to proofread, suggest synonyms, check grammar and spelling, and debug R script. All suggestions made by these tools have been scrutinized and cross-checked by the authors.

To prolong the thrill of life cycles, you could play at surviving the winter like a sexual lineage of poplar rust. https://itch.io/jam/scientific-game-jam-100-en-ligne/rate/791842

## Conflicts of Interest

The authors declare no conflicts of interest

## Benefit-Sharing Statement

The advantage of this study is that its analysis code is an automated workflow and data are publicly accessible.

## Supplementary figures

**Figure S1.**
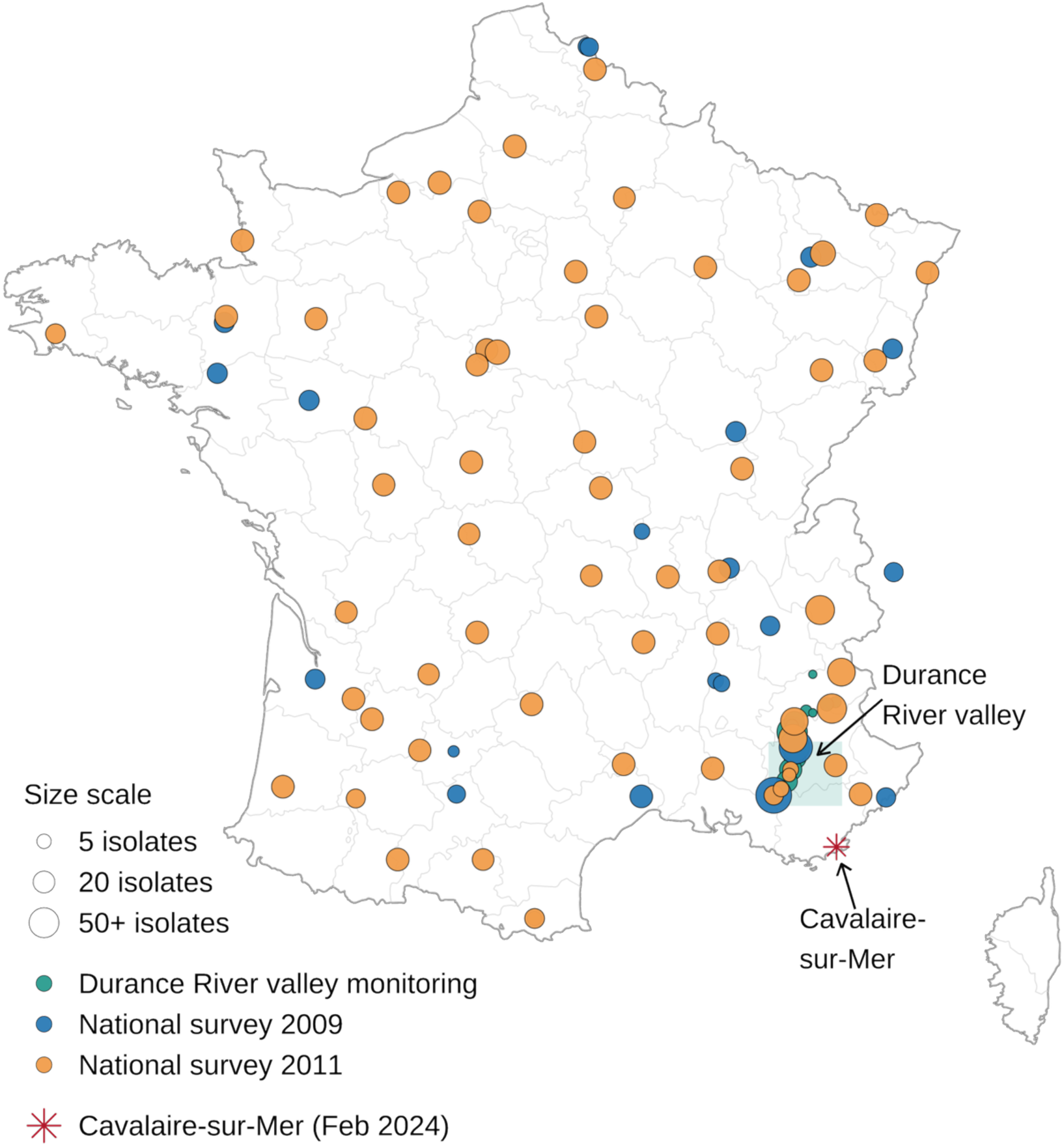
Geographic distribution of *Melampsora larici-populina* individuals collected across France. Each circle represents a sampling site, with size proportional to the number of individuals collected (5, 20, or 50+ individuals). Colors indicate the sampling campaign: blue for the 2009 national survey, orange for the 2011 national survey (n = 60 sites), and teal for the long-term monitoring program in the Durance River valley (surveys conducted in 2004, 2008, and 2020). The red asterisk marks the collection at Cavalaire-sur-Mer in February 2014 and 2024. Individuals collected before 2004 are not shown on this map but were included in the population genetic analyses.

**Figure S2.**
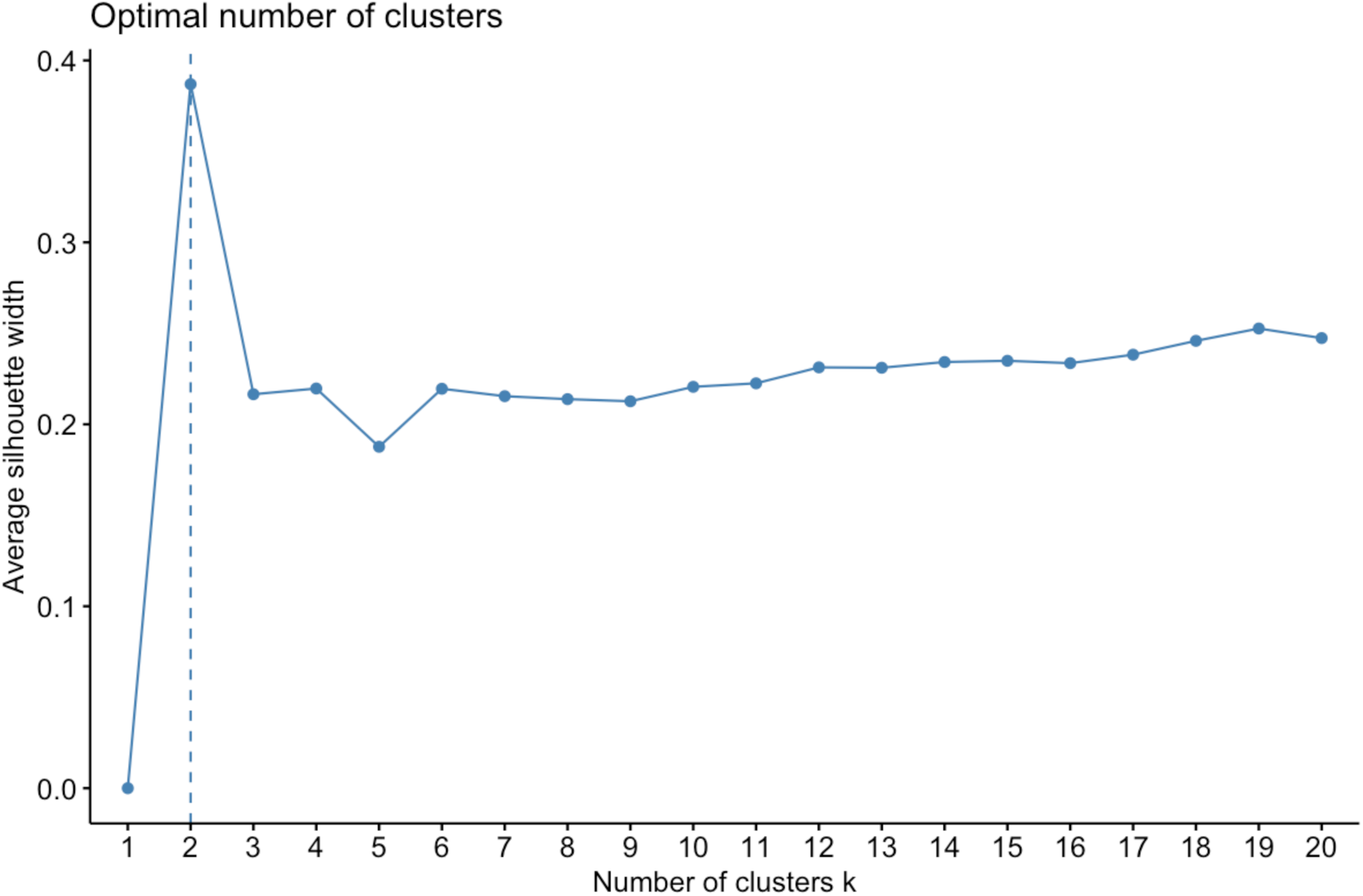
Silhouette analysis for determining the optimal number of genetic clusters (k) in *Melampsora larici-populina*. Average silhouette width was computed for k = 1 to 20 using K-means clustering applied to the first 10 principal components of the microsatellite allele frequency matrix. The dashed vertical line indicates the optimal k = 2, which achieved the highest average silhouette width (≈ 0.4), indicating the most likely distinct and well-separated cluster structure.

**Figure S3.**
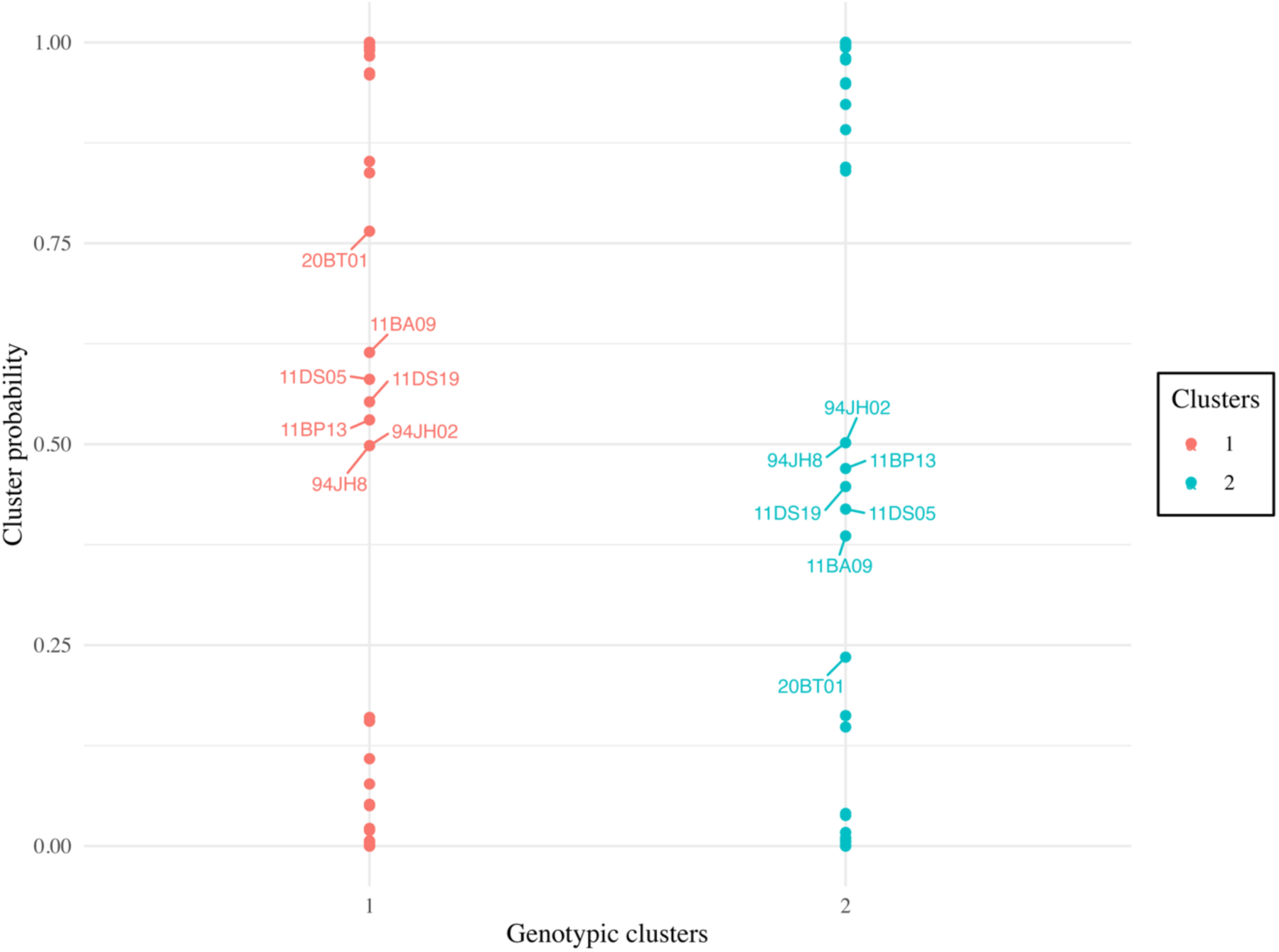
Scatter plot illustrating the membership probability for individuals within each genotypic cluster, with the names of individuals of lower probability.

**Figure S4.**
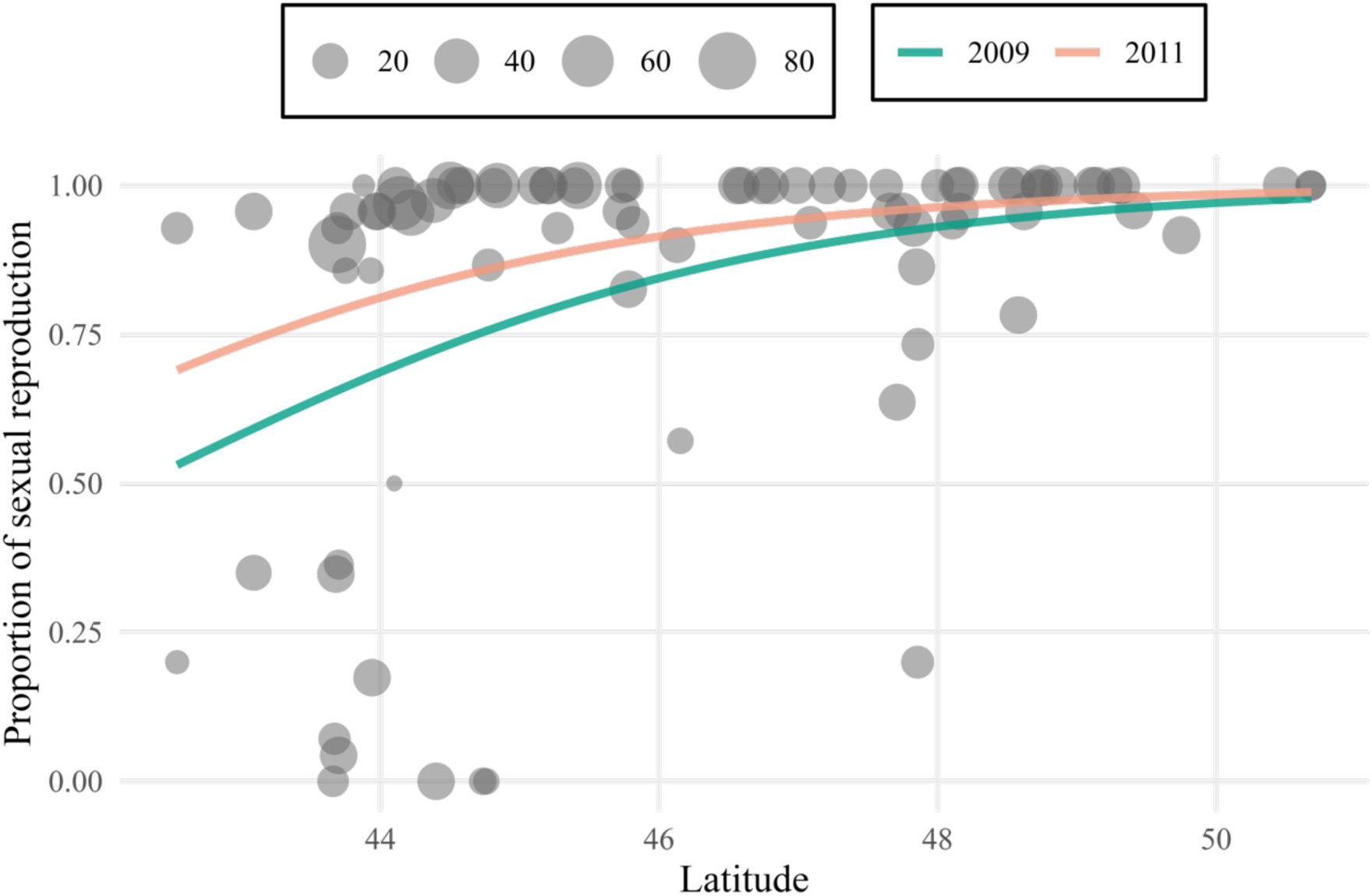
The regression plot illustrates the relationship between latitude and the proportion of sexual reproduction. This relationship was analysed using a binomial generalized linear mixed model (GLMM). It was based on data from 2009 and 2011. The size of the points represents sample sizes.

## Supplementary Tables

**Table S1.** The data set for 2,122 *Melampsora larici-populina* individuals used in population genetic analyses includes collection year, geographic location, and genotypes across 21 microsatellite markers.

**Table S2.** Detailed information on samples collected at Cavalaire-sur-Mer, with their reproductive mode, sample size, and the years in which each MLL was detected.

| Year | MLL | Reproduction | N | MLG | Years sampled overall |
|---|---|---|---|---|---|
| 2014 | 2 | Asex | 6 | 1 | 2014, 2020 |
| 2014 | 1 | Asex | 1 | 1 | 1992, 2004, 2008, 2009, 2011, 2014, 2020, 2024 |
| 2014 | 8 | Asex | 1 | 1 | 1993, 1996, 2004, 2011, 2014 |
| 2024 | 5 | Sex | 26 | 1 | 2024 |
| 2024 | 35 | Asex | 4 | 1 | 2024 |
| 2024 | 56 | Sex | 3 | 1 | 2024 |
| 2024 | 1 | Asex | 2 | 1 | 1992, 2004, 2008, 2009, 2011, 2014, 2020, 2024 |
| 2024 | 120 | Sex | 2 | 2 | 2024 |
| 2024 | 1383 | Sex | 1 | 1 | 2024 |
| 2024 | 1384 | Sex | 1 | 1 | 2024 |
| 2024 | 1385 | Sex | 1 | 1 | 2024 |

For each multilocus lineage (MLL) collected at Cavalaire-sur-Mer, the table gives the collection year, reproductive mode, number of isolates (N), number of distinct multilocus genotypes (MLG), and all years in which that lineage was detected in the dataset.

## Notes

https://doi.org/10.5281/zenodo.15100450

### Competing Interest Statement

The authors have declared no competing interest.

### Summary of Updates

New updates according to the PCI recommender

